# Evidence for a putative isoprene reductase in *Acetobacterium wieringae*

**DOI:** 10.1101/2022.11.16.516518

**Authors:** Miriam Kronen, Xabier Vázquez-Campos, Marc R. Wilkins, Matthew Lee, Michael J Manefield

## Abstract

Recent discoveries of isoprene-metabolizing microorganisms suggest they might play an important role in the global isoprene budget. Under anoxic conditions, isoprene can be used as an electron acceptor and is reduced to methylbutene. This study describes the proteogenomic profiling of an isoprene-reducing bacterial culture to identify organisms and genes responsible for the isoprene hydrogenation reaction. A metagenome assembled genome (MAG) of the most abundant (89% rel. abundance) lineage in the enrichment, *Acetobacterium wieringae*, was obtained. Comparative proteogenomics and RT-PCR identified a putative five-gene operon from the *A. wieringae* MAG upregulated during isoprene reduction. The operon encodes a putative oxidoreductase, three pleiotropic nickel chaperones (2 x HypA-like, HypB-like) and one 4Fe-4S ferredoxin. The oxidoreductase is proposed as the putative isoprene reductase with a binding site for NADH, FAD and two pairs of [4Fe-4S]-clusters. Other known *Acetobacterium* strains do not encode the isoprene-regulated operon but encode, like many other bacteria, a homolog of the putative isoprene reductase (∼47–49% amino acid sequence identity). Uncharacterized homologs of the putative isoprene reductase are observed across the *Firmicutes, Spirochaetes, Tenericutes, Actinobacteria, Chloroflexi, Bacteroidetes* and *Proteobacteria*, suggesting the ability of biohydrogenation of unfunctionalized conjugated doubled bonds in other unsaturated hydrocarbons.

**Importance:** Isoprene was recently shown to act as an electron acceptor for a homoacetogenic bacterium. The focus of this study is the molecular basis for isoprene reduction. By comparing a genome from our isoprene reducing enrichment culture, dominated by *Acetobacterium wieringae*, with genomes of other *Acetobacterium* lineages that do not reduce isoprene, we shortlisted candidate genes for isoprene reduction. Using comparative proteogenomics and reverse transcription PCR we have identified a putative five-gene operon encoding an oxidoreductase referred to as putative isoprene reductase.

## Introduction

Isoprene represents the most abundant biogenic volatile organic compound (BVOC) on Earth and accounts for 70% of total annual BVOC emissions excluding methane (1–4). Large quantities of isoprene (500 to 600 Tg per year; (5–7)) enter the atmosphere making it an important participant in atmospheric chemistry, contributing to ozone and secondary organic aerosol (SOA) formation in the troposphere and increasing the lifetime of methane indirectly (8–13). It is mainly produced by plants but also by other organisms such as Gram-positive and Gram-negative bacteria (14–18), fungi (19) and algae (20, 21).

Soils and marine environments harbouring aerobic isoprene degrading organisms serve as isoprene sinks (4, 22). The fate of isoprene under anoxic conditions was examined recently (23), whereby isoprene was found to act as a 2e^-^ acceptor with one C=C bond being reduced by an anaerobic enrichment culture to predominately 2-methyl-1-butene. Sequencing of 16S rRNA gene amplicons from this culture revealed enrichment of *Acetobacterium* to 92–100% relative abundance with *Comamonadaceae* accounting for the rest (2–8%). The homoacetogenic *Acetobacterium* lineage dominating the H_2_ fed enrichment utilized 1.6 μmol isoprene h^-1^ as an electron acceptor in addition to HCO_3_^−^. Growth of the homoacetogen with isoprene produces 40% less acetate than with H_2_ and HCO_3_^−^ alone suggesting its reduction to methylbutene is coupled to energy conservation (23).

Homoacetogens are known to reduce electron acceptors other than CO_2_, such as fumarate (24), nitrate (25), chloroethenes, chloroethanes (26), brominated aromatics (27) and acrylate derivatives (28). Reduction of the functionalized C=C bond (C=C bond conjugated to an electron-withdrawing group) in caffeate by the model organism *Acetobacterium woodii* (28– 30) is a well-studied example for CO_2_-alternative electron acceptors in *Acetobacterium* species. The caffeate reduction operon and corresponding enzymes have been characterised in *A. woodii* (30, 31). A caffeyl-CoA reductase-Etf complex uses flavin-dependent electron bifurcation to drive the endergonic reduction of ferredoxin with NADH as electron donor by coupling it to the exergonic NADH-dependent reduction of caffeyl-CoA (32).

This study aimed to identify microorganisms and their corresponding genes/enzymes involved in the reduction of the unfunctionalized C=C bond in isoprene. DNA from an isoprene-reducing enrichment culture was sequenced and protein profiles with and without isoprene were compared. Metagenomic and comparative metaproteomic analyses implicate a putative oxidoreductase in isoprene reduction encoded in a putative five-gene operon.

## Results

### A putative oxidoreductase encoded by *Acetobacterium wieringa*e is upregulated by isoprene

Cell suspension experiments with the *Acetobacterium*-dominated (rel. abundance 16S rRNA gene amplicon sequencing 92–100%) enrichment culture (23) pre-grown on H_2_/HCO_3_^-^ showed that isoprene reduction is induced in the presence of isoprene, H_2_ and HCO_3_^-^ (See SI for details; **Figure S1**).Therefore, label-free comparative metaproteomics was performed to identify proteins and corresponding genes involved in isoprene metabolism. To generate a database for protein identification, the isoprene reducing enrichment culture was grown on H_2_/HCO_3_^-^ /(±isoprene), and the extracted DNA was sequenced. Over 7.5 million paired-end reads were used to assemble 338 contigs. Two near-complete metagenome-assembled genomes (MAGs) were obtained (**Table 1**). No other lineages were detected. MAG ISORED-1 showed 74% average amino acid identity (AAI) and 79% average nucleotide identity (ANI) to *Comamonas aquatica* CJD (**Table S1**) and ISORED-2 showed 97% AAI and ANI to *Acetobacterium wieringae* DSM 1911 (**Table S2**). MAG ISORED-2 is dominant in the enrichment based on relative abundance (77%) and more so when cultivated in the presence of isoprene (rel. abundance 88.71% metagenome) (**Table 1**).

**Table 1.** Summary of metagenome assembled genomes from isoprene reducing culture. Biomass contribution was calculated according to (99).

| MAGS | Genome size (Mbp) | GC content (%) | Completeness / Redundancy (%) | Mean Coverage | Relative abundance (% metagenome) |  | Relative abundance (% biomass) |  |
| --- | --- | --- | --- | --- | --- | --- | --- | --- |
|  |  |  |  |  | H<sub>2</sub>/HCO<sub>3</sub><sup>-</sup> | Isoprene | H<sub>2</sub>/HCO<sub>3</sub><sup>-</sup> | Isoprene |
| *Comamonas* sp.<br>ISORED-1<br>(GCA_902175065.1) | 3.22 | 63.7 | 98.28–99.3 / 0.72–3.45 | 50.8 x | 22.99 | 11.29 | 15.54 | 6.41 |
| *Acetobacterium wieringae*<br>ISORED-2<br>(GCA_902175055.1) | 3.81 | 44 | 92.93–99.28 / 0.95–5.76 | 244.3 x | 77.01 | 88.71 | 84.46 | 93.59 |

Differential metaproteomes were generated by LC-MS/MS analysis of proteins obtained from cultures grown on H_2_/HCO_3_^-^ with and without isoprene. A total of 1531 proteins were identified. Consistent with the dominance of the homoacetogen in culture, 1279 proteins (83.5%) belonged to *A. wieringae* ISORED-2 (**Figure 1B)** and 252 proteins (16.5%) were assigned to *Comamonas* ISORED-1 (**Figure 1A**). This is also reflected in the estimated biomass contributions where *A. wieringae* ISORED-2 is constituting ∼94% and *Comamonas* ISORED-1 ∼6% rel. abundance of biomass of the community (**Table 1**).

**Figure 1.**
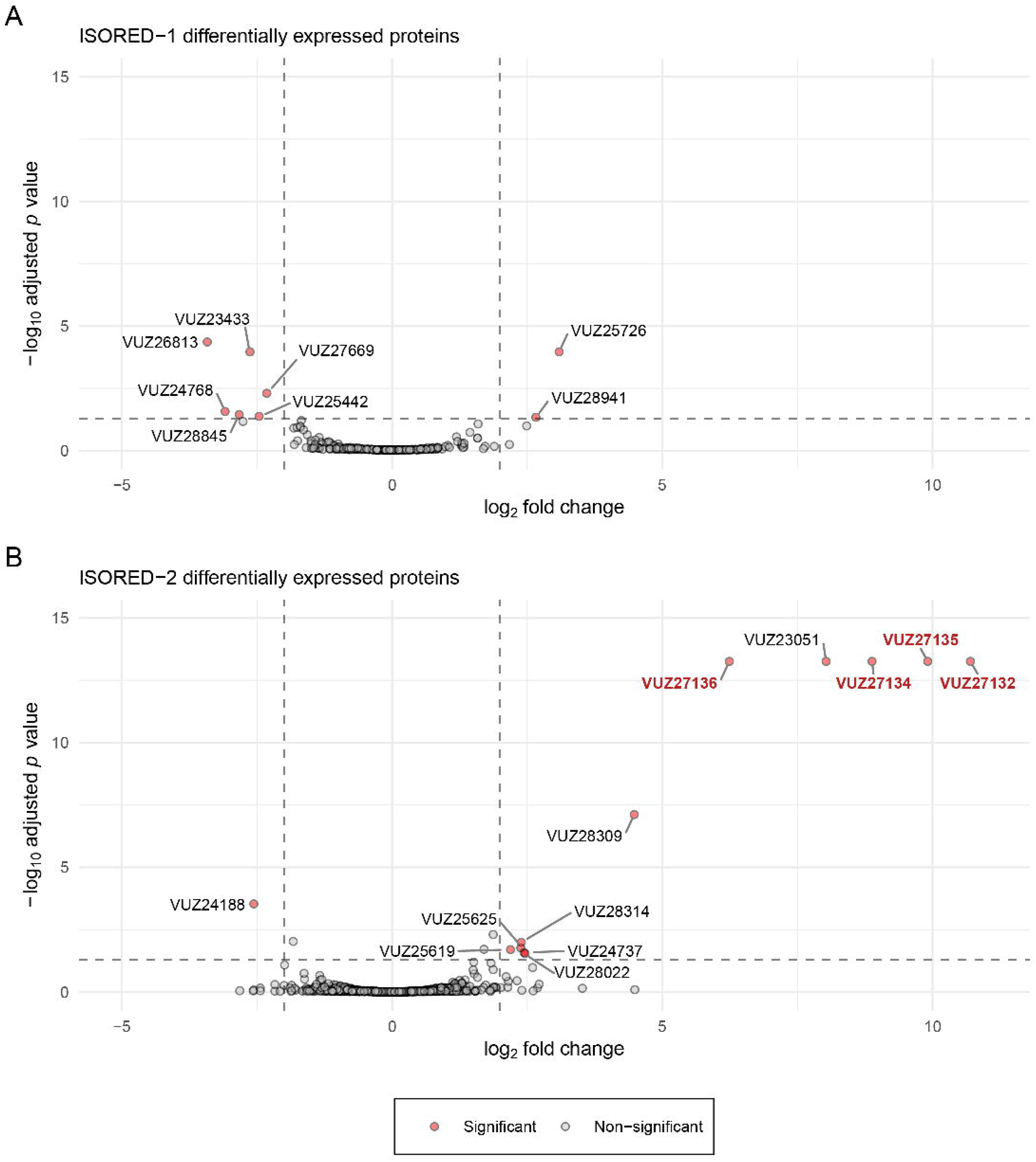
Volcano plot of the metaproteomic data comparing cells grown on H_2_/HCO_3_^-^/isoprene vs. H_2_/HCO_3_^-^. Significant data points (coloured) are based on a LFC of ±2 and an adjusted *p*-value of ≤0.05. Labelling of the significant points is based on metagenome assembled genomes ISORED-1 (A, *Comamonas* sp.) and ISORED-2 (B, *Acetobacterium wieringae*). Proteins located adjacent to each other in the genome of *A. wieringae* ISORED-2 (B) are highlighted. Data was obtained from 4 replicates for each growth condition (PRIDE database PXD023683).

Differential expression analysis identified changes in 12 proteins from *A. wieringae* ISORED-2 and 8 proteins of *Comamonas* sp. ISORED-1 between cells grown on H_2_/HCO_3_^-^/isoprene vs. H_2_/HCO_3_^-^ (**Table 2**). Only 13 proteins were upregulated in isoprene-exposed cells, 11 belonging to *A. wieringae* ISORED-2 and 2 belonging to *Comamonas* sp. ISORED-1 (**Figure 1AB, Table 2**).

**Table 2.**
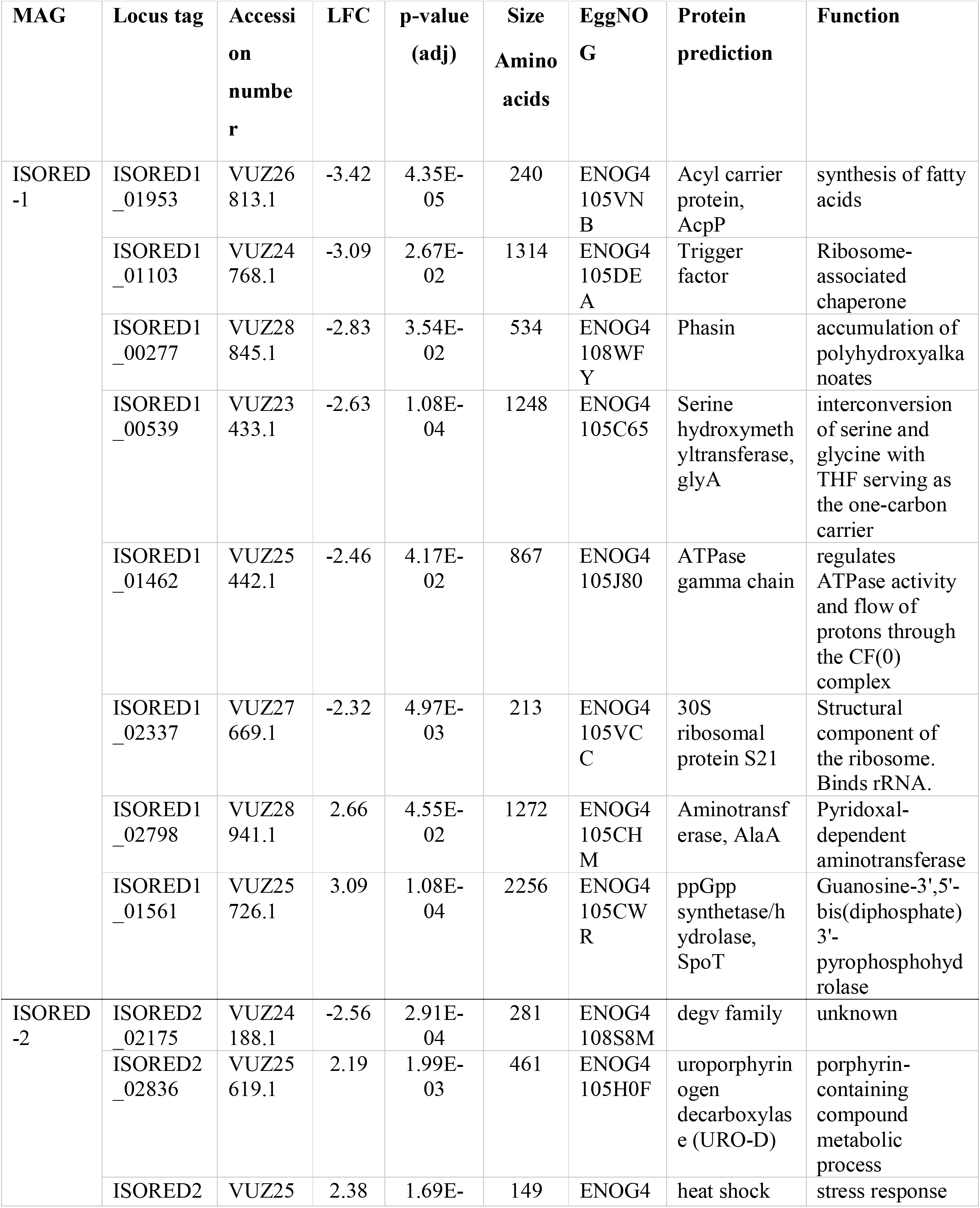
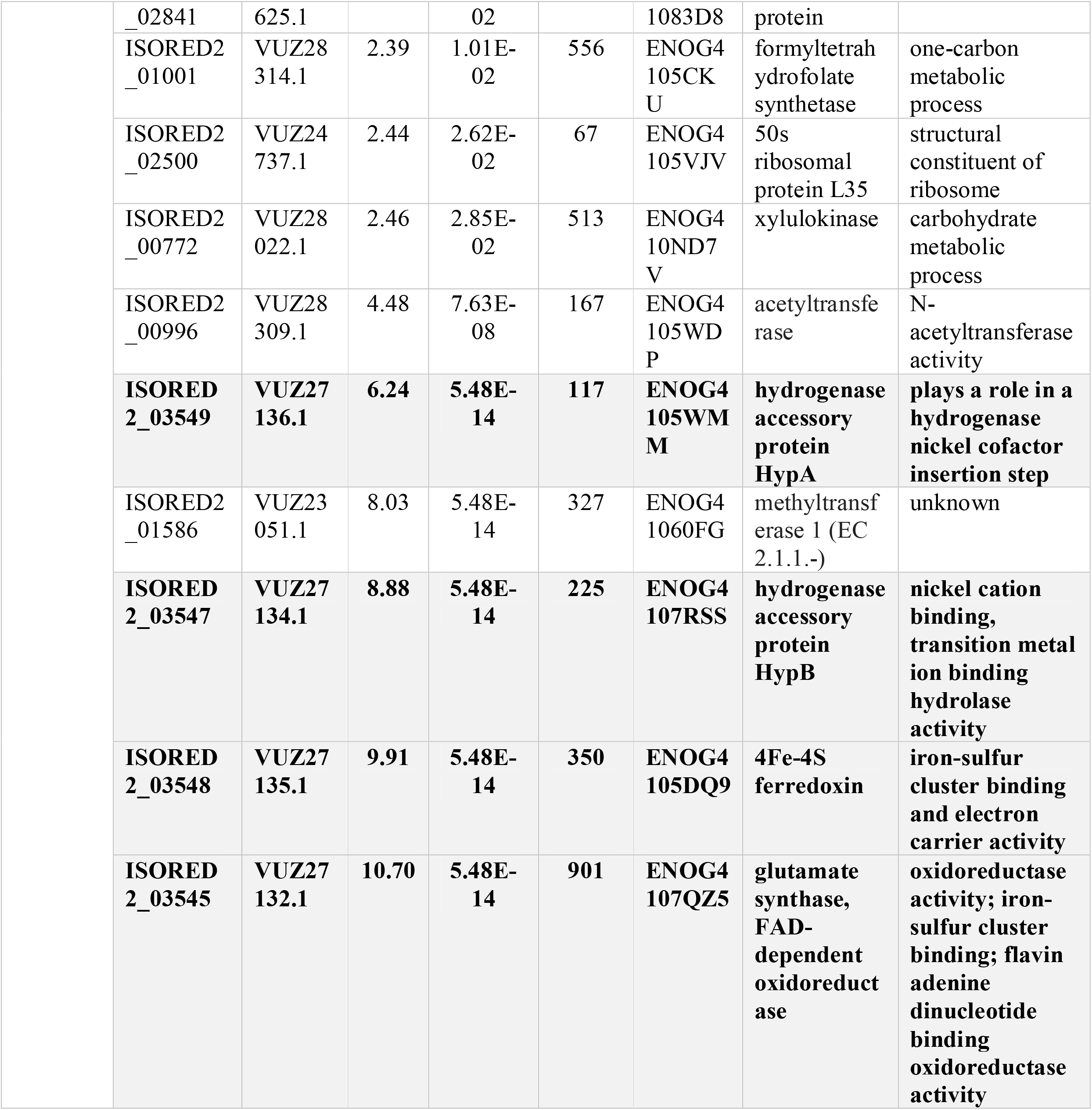
List of proteins that significantly differed in abundance between cells grown on H_2_/HCO_3_^-^/isoprene vs. H_2_/HCO_3_ ^-^ in the ISORED-2 (*A. wieringae*) metagenome assembled genome. Significant data points are based on a minimum abs(logFC) of 2 and an adjusted p-value of 0.05. EggNOG matches are shown with their functional description. Proteins encoded by genes located adjacent to each other are highlighted in bold and grey. Data was obtained from 4 replicates for each growth condition (PRIDE database (PXD023683)).

Five out of 12 isoprene-responsive proteins from *A. wieringae* were more highly upregulated, with a log2 fold change (LFC) of 6.2–10.7, compared to the remaining six (LFC 2.5–4.8), and one protein was downregulated (VUZ24188.1, LFC -2.56). Four (VUZ27132.1, VUZ27134.1, VUZ27135.1, VUZ27136.1) out of these five are adjacent to each other in the ISORED-2 MAG indicating they might belong to the same operon (**Figure 1B, Figure 2A, Table 2**). Protein VUZ27132.1 (**Table 2**) is predicted to be a molybdopterin oxidoreductase with the best orthologous group match ENOG4107QZ5. Protein VUZ27136.1 (ENOG4105WMM) is a HypA-like homolog, and VUZ27134.1 (ENOG4107RSS) is a HypB-like homolog. HypAB proteins are typically responsible for the acquisition and insertion of nickel in the catalytic centre of [NiFe]-hydrogenases. Protein VUZ27135.1 (ENOG4105DQ9) belongs to the 4Fe-4S superfamily (SSF54862), but the sequence is not affiliated with a specific family. Protein VUZ23051.1 (ENOG41060FG, methyltransferase) is also highly expressed (LFC 8.03) but is not part of the operon. Predicted protein functions from the six less isoprene-responsive proteins (LFC 2.5–4.8) include acetyltransferase, xylulokinase, uroporphyrinogen decarboxylase, heat shock protein, formyltetrahydrofolate synthetase and 50S ribosomal protein L35 (**Table 2**).

**Figure 2.**
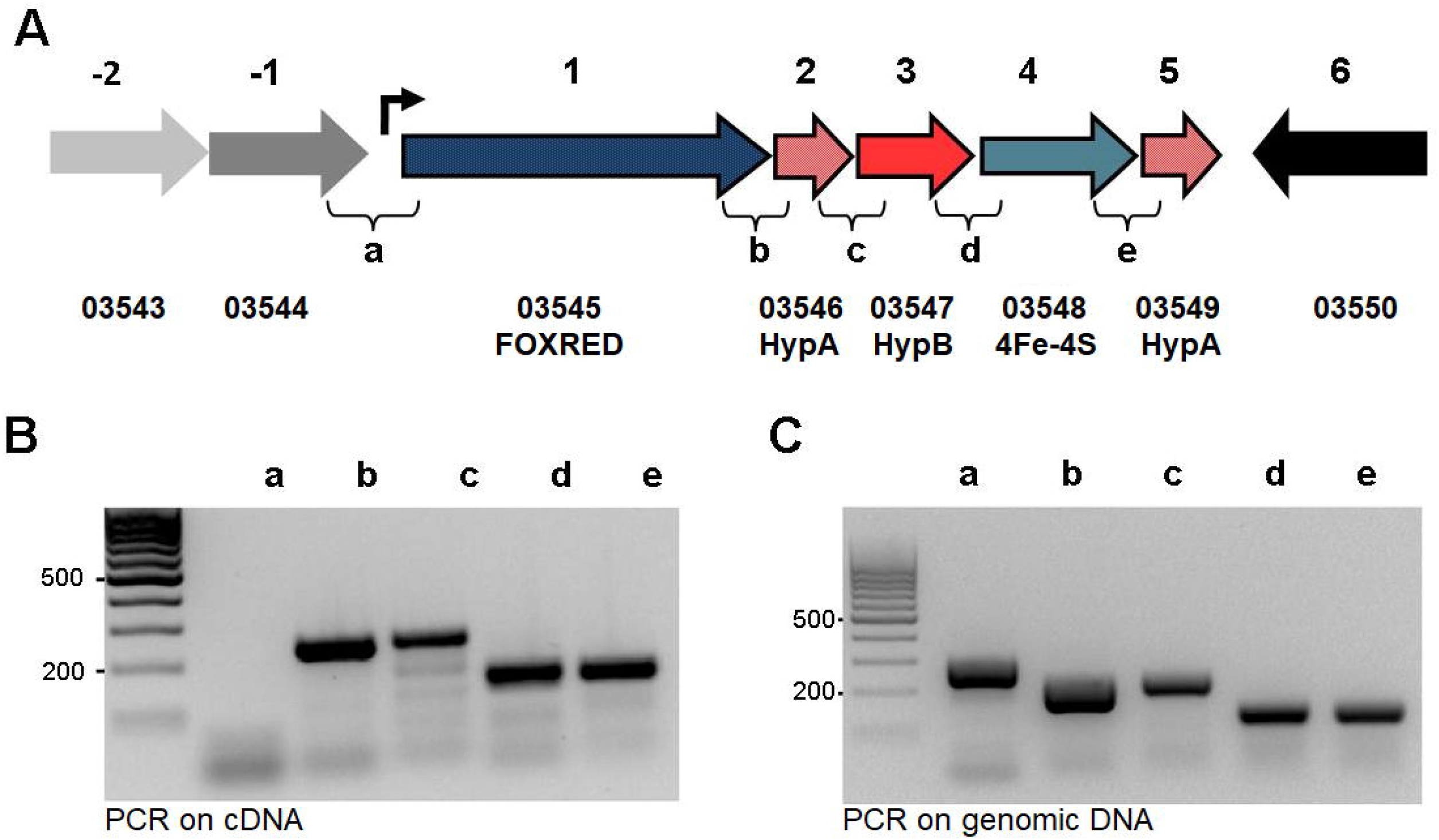
Organisation of genes upregulated by isoprene and confirmation of operon structure by RT-PCR. (**A**) Five gene operon harbours genes encoding putative FAD dependent oxidoreductase (FoxRed, ISORED2_03545, blue), three nickel-inserting, hydrogenase maturation factors HypA (ISORED2_03546, ISORED2_03549, light red), HypB (ISORED2_03547, dark red) and one 4Fe-4S ferredoxin (ISORED2_03548, turquoise), diagrammatic representation of the operon with open reading frames (ORF) 1–5 and their intersections annotated a–e; (**B**) amplicons with primers connecting intersections of the neighbouring ORFs on cDNA transcript; and (**C**) amplicons with primers connecting intersections of the neighbouring ORFs on chromosomal DNA positive control. No bands appeared in negative controls lacking reverse transcriptase.

In the *Comamonas* sp. lineage two proteins were upregulated upon isoprene exposure; protein VUZ25726.1 (ENOG4105CWR) (LFC 3.09) a (p)ppGpp synthetase/hydrolase (SpoT) and protein VUZ28941.1 (ENOG4105CHM) (LFC 2.66) an aminotransferase (AlaA) (**Table 2**). SpoT is typically involved in the stringent response (33), a ubiquitous stress signalling pathway that enables bacteria to respond to nutrient starvation and AlaA catalyses the reversible transamination reaction pyruvate + glutamate ↔ L-alanine + α-ketoglutaric acid. *Comamonas* sp. ISORED-1 had 6 proteins downregulated (LFC -3.42 – -2.31) upon isoprene exposure; an acyl carrier protein (AcpP), a trigger factor (ribosome associated chaperon), phasin (involved in accumulation of polyhydroxyalkanoates), serine hydroxymethyltransferase (glyA), ATPase gamma chain and 30S ribosomal protein S21 (**Table 2**).

### The putative isoprene operon

Apart from *A*.*wieringae* ISORED-2’s oxidoreductase (VUZ27132.1), no isoprene responsive protein from MAG ISORED-2 or ISORED-1 is predicted by protein function to be involved in redox processes (**Table 2**). This makes the oxidoreductase from the *A. wieringae* lineage, the only likely candidate that could catalyse the isoprene hydrogenation reaction. The relevant gene of *A. wieringae*’s oxidoreductase (VUZ27132.1) is organised in one gene cluster together with the genes of three other isoprene-induced proteins (VUZ27134.1, VUZ27135.1, VUZ27136.1) and an additional gene (encoding HypA-like protein, VUZ27133.1) not upregulated in the metaproteome (**Figure 2A**). All five genes have the same orientation, with intergenic regions ranging between 7–71 nucleotides. Operon prediction analysis (FGENESB) suggested the five genes are transcribed as an operon (**Figure 2A, Figure S2**). Reverse transcription PCR using primer sets flanking individual intergenic regions of adjacent genes (**Table S3 and S4**) also indicated that the genes are transcribed as an operon (**Figure 2BC**). For promoter-prediction analysis (BPROM) see SI text (**Figure S2**).

### Isoprene reduction and the putative isoprene-regulated operon is unique to ISORED-2 amongst known *Acetobacterium* species

Whilst *A. wieringae* DSM 1911 is the same species as *A. wieringae* ISORED-2, it did not catalyse isoprene reduction nor did *A. woodii* DSM 1030, *A. malicum* DSM 4132 or *A. dehalogenans* DSM 11527 (23). Comparative pangenome analysis with eight publicly available *Acetobacterium* genomes (**Table S5**) was used to assess the distribution of the putative isoprene-regulated operon encoding the oxidoreductase (VUZ27132.1) within the *Acetobacterium* genus and to find features unique to the *A. wieringae* ISORED-2 MAG.

Pangenome analysis showed the protein coding sequences from all nine genomes (33035 in total) grouped into 8190 gene clusters, based on an MCL inflation value of 6 (parameter controlling the granularity of the clustering) (**Figure S3**). A shared set of 1492 gene clusters (core) is shown across the nine genomes along with protein sets that are unique in each of the *Acetobacterium* genomes (**Figure S3, Table S6)**.

For the *A. wieringae* ISORED-2 MAG, a total repertoire of 3628 proteins in 3386 gene clusters were identified. Of these gene clusters, 318 with 352 proteins were unique to the *A. wieringae* ISORED-2 MAG (**Table S7**). Regardless of the MCL inflation value (2,4,6) used, the oxidoreductase (VUZ27132.1) from the putative isoprene-regulated operon was always located in a singleton gene cluster. Amongst *Acetobacterium* genomes, four genes in the putative isoprene-regulated operon are unique to the ISORED-2 MAG; 4Fe-4S ferredoxin, two HypA-like proteins and the oxidoreductase (VUZ27132.1). This is consistent with the absence of isoprene reducing ability in *A. woodii* DSM 1030, *A. wieringae* DSM 1911, *A. malicum* DSM 4132 and *A. dehalogenans* DSM 11527.

The putative isoprene-regulated operon (hereafter referred to as *isr* operon) is located between 69745–75048 bp in a 90374 bp contig (**Figure 3B**). The first half of this contig contains mainly protein-coding genes of viral origin. The mean contig coverage and the coverage of the proviral portion are close to the values for the MAG indicating no active viral replication. The provirus (*Siphoviridae*) shows an average amino acid identity of 59.11% with the *Erysipelothrix* phage Φ1605 (34) based on CheckV, and tBLASTx of many of the viral proteins also show similar identities with several *Streptococcus* phages (35). This proviral region also appears in other *Acetobacterium* spp. genomes including *Acetobacterium wieringae* DSM 1911 and *Acetobacterium* sp. KB-1 (**Figure 3AC**). The contig contains three different Ser-recombinases (integrases). Two of them adjacent to the provirus (**Figure 3B**), show very high identity values with recombinases found in *A. wieringae* DSM 1911, *A*. sp. MES1, and *A*. sp. KB-1 and might be part of the provirus.

**Figure 3.**
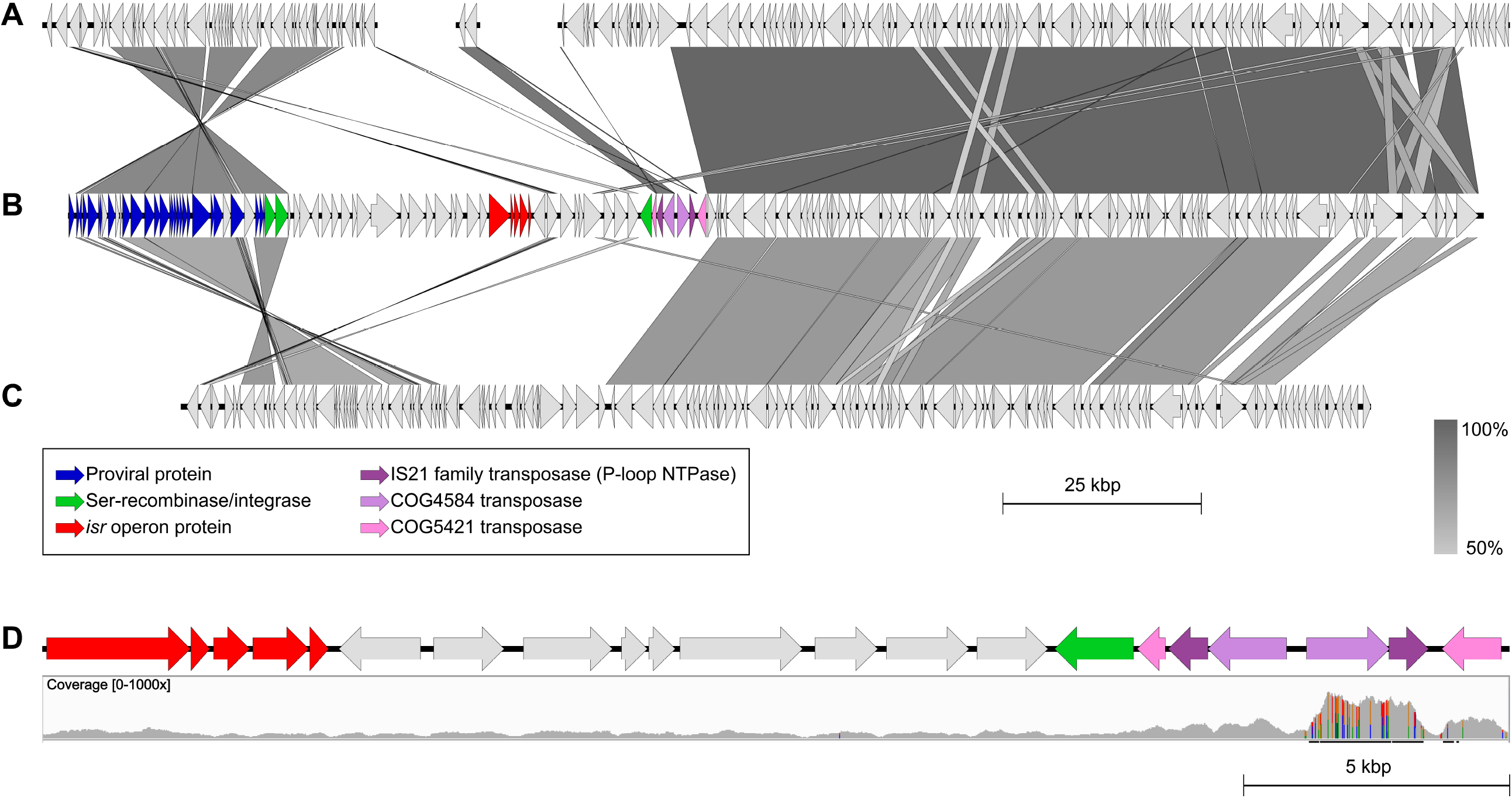
Details of the reassembled contig containing the putative isoprene reductase operon. **A-C**) Homology between *A. wieringae* DSM 1911 (**A**, contigs LKEU01000044, 59 and 37), *A. wieringae* ISORED-2 (**B**) and *A*. sp. KB-1 (**C**, between 925–1,075 kbp). Grayscale gradient indicates the percentage identity (Easyfig blastn min. size 100, e-value 1e-5, min. identity 50%). **D**) Close view at the coordinates between the isoprene-regulated operon and the IS elements highlighting the coverage of the region (max. 748x).

As the provirus and recombinases were the main sequences sharing some degree of synteny in the original contig with assembled *Acetobacterium* spp. genomes, the contig encoding the putative *isr* operon was scrutinised in more detail and the MAG reassembled to better understand the gene environment of the operon (see SI). The reassembled ISORED-2 MAG was ∼40 kbp larger with a more polished putative *isr* operon contig (**Figure 3BD**), which showed a higher degree of synteny with the genome of *Acetobacterium* sp. KB-1 (**Figure 3C**), the most complete *Acetobacterium* spp. genome to date. While the reassembly process was able to extend the putative *isr* operon contig, it did so at the expense of collapsing several insertion sequences (IS). Most of the IS in this region are suspected to appear in tandem repeats based on their higher apparent sequence coverage.

### Taxonomic distribution and phylogeny of isoprene-induced oxidoreductase homologs

We searched for *A. wieringae*’s oxidoreductase (VUZ27132.1) within the EggNOG and NCBI databases to determine its broader distribution and to find biochemically characterized homologs. Protein VUZ27132.1, which is 901 amino acids long, is predicted to belong to the bacterial orthologous group ENOG4107QZ5 (141 proteins, 99 species), with an archaeal counterpart in the orthologous group arCOG01292 (88 proteins, 55 species). The ENOG4107QZ5/arCOG01292 orthologous groups primarily (92%) contain proteins of ∼500– 600 amino acids long whereas the NCBI BLAST search primarily (95%) gave results of full-length homologs ∼900 amino acids (**Table S8, Figure 4A**). All sequences share a homologous domain of 500 amino acids that aligns to position 300-800 amino acids in protein VUZ27132.1.

**Figure 4.**
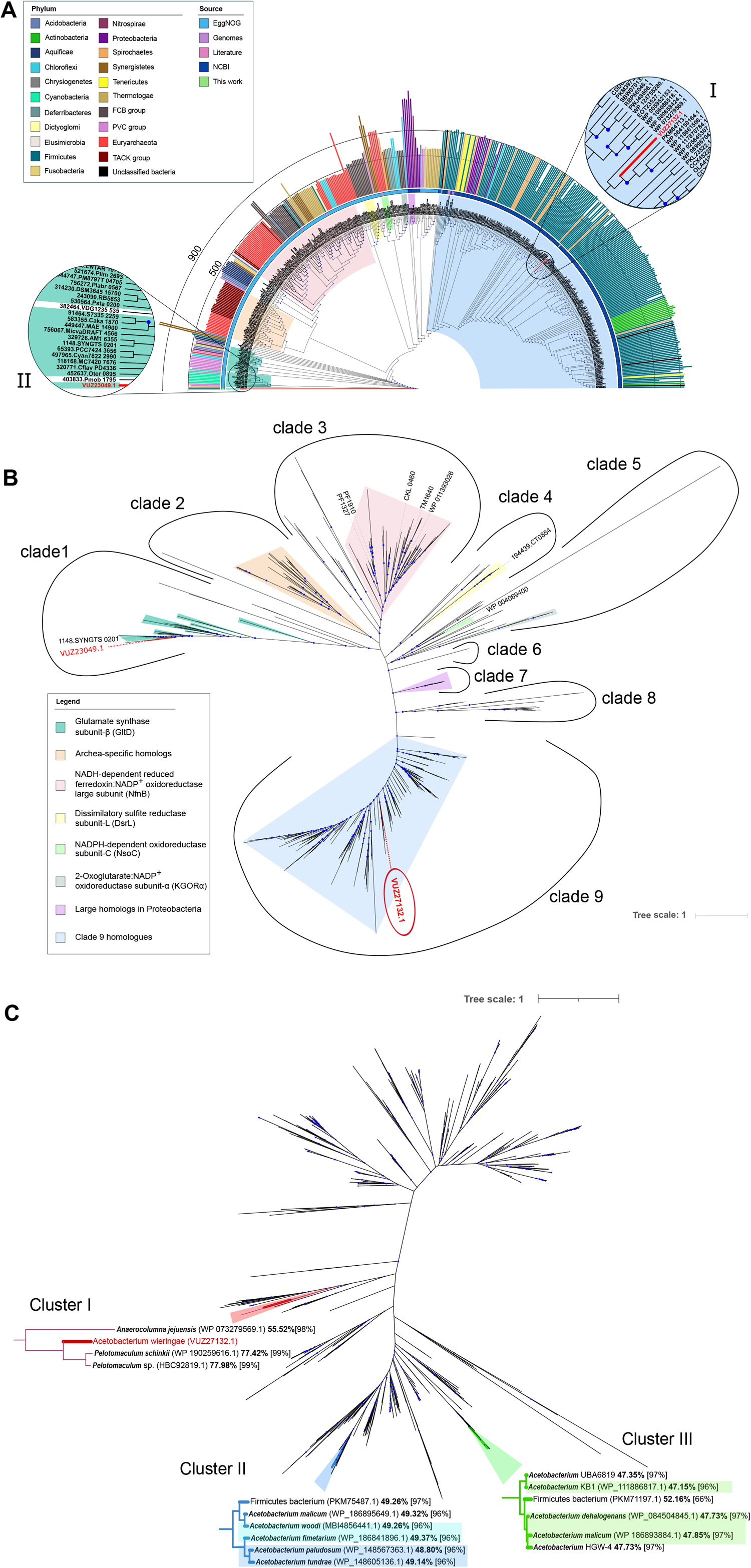
A Unrooted maximum likelihood phylogenetic tree of IsrA (VUZ27132.1) and its homologs. Phylogeny of amino acid sequences of IsrA and its homologs drawn as an unrooted circular tree. Highlighted are the positions of protein VUZ27132.1 (I) and another VUZ27132.1 ortholog from *A. wieringae* ISORED-2 VUZ23049.1 (II). The outer circle shows bars representing the protein length coloured according to phylum. Outer ring indicates the source of the sequences. Clades are coloured according to the clades in the unrooted tree (**Figure 4B**). **4B)** The phylogenetic analysis of IsrA and all its homologs revealed nine distinct clades containing redox enzymes involved in different reactions. Blue circles indicate an ultrafast bootstrap support ≥90%. Proteins labelled in black indicate characterized proteins. See **Table S7** for information about each sequence. See also interactive iTol link https://itol.embl.de/tree/4918010318257311557452600. **4C)** Extended phylogenetic analysis of clade 9 homologs. Subtrees show clusters of IsrA homologs from *Acetobacterium* spp. and the closest homologs to IsrA (*Pelotomaculum* spp.). Blue circles indicate an ultrafast bootstrap support ≥90%. Amino acid identity values are shown next to the name followed by sequence coverage between brackets. Highlighted *Acetobacterium* names indicate identical gene arrangement (refer to **Figure S4**). See **Table S8** for information about each sequence.

Phylogenetic analysis of protein VUZ27132.1 and its homologs revealed nine distinct clades (**Figure 4B)**. Some sequences are grouped into main lineages located within the clades by analysis of the genomic context and/or information from the literature about the protein’s enzymatic activity: (1) Glutamate synthase β-subunit (GltD) homologs (36), (2) *Archaea*-specific homologs, (3) NADH-dependent reduced ferredoxin:NADP^+^-oxidoreductase large subunit (NfnB) homologs (37–40), (4) Dissimilatory sulfite reductase subunit L (DsrL) homologs(41–45), (5) NADPH-dependent oxidoreductase subunit C (NsoC) homologs (46) and 2-Oxoglutarate:NADP^+^-oxidoreductase subunit α (KGOR α) (47), (7) large homologs in *Proteobacteria* and (9) clade 9 homologs (**Figure 4B, Table S8**). MAG ISORED-2 encodes another member of the ENOG4107QZ5 orthologous group (protein VUZ23049.1) but it clusters with the glutamate synthase β-subunit (GltD) sequences in clade 1 (**Figure 4A II**). The families *Desulfobacteraceae* and *Syntrophobacteraceae* contain an even longer ortholog (∼1300–1400 amino acids) (clade 7, **Figure 4AB)**.

The clade 9 homologs are defined as the most closely related to *A. wieringae*’s isoprene-upregulated oxidoreductase (VUZ27132.1); they all aligned to the entire 901 amino acids sequence. They are mainly distributed among *Firmicutes* strains, but some are found in *Spirochaetes, Tenericutes, Actinobacteria* as well as *Chloroflexi, Bacteroidetes* and *Proteobacteria* (**Figure 4A, Table S8**). Most taxa containing clade 9 homologs are strict anaerobes (detailed information in Supplement Material, **Table S8**).

To determine uniqueness and conservation of protein VUZ27132.1 and the putative *isr* operon, a more detailed and extended phylogenetic tree of clade 9 sequences was generated. Other *Acetobacterium* strains also contain clade 9 proteins which share ∼47–49% amino acid sequence identity with protein VUZ27132.1 (**Figure 4C, Figure S4, Table S9**). Protein VUZ27132.1 from *A. wieringae* ISORED-2 is distinct from any other *Acetobacterium* homolog in a subclade (**Figure 4C, cluster I**) while homologs from *A. woodii, A. fimetarium, A. malicum* (WP_186895649.1), *A. paludosum*, and *A. tundrae* are clustered together in a separate subclade (**Figure 4C, cluster II**). Homologs from *A. dehalogenans* DSM 11527, *A*. sp. HGW-4, *A*. sp. UBA6819, *A. malicum* (WP 186893884.1) and *A*. sp. KB-1 are located in a third subclade (**Figure 4C, cluster III**). Besides sharing proximity to *hypA* and *hypB*, the gene neighbourhoods of clade 9 proteins in *Acetobacterium* spp. from clusters II and III are distinct from that of *A. wieringae* ISORED-2 (**Figure S5**). In case of cluster III (**Figure 4C)**, the oxidoreductase gene is flanked by another ferredoxin oxidoreductase (ENOG41061TH) and a helix-turn-helix domain containing protein (ENOG4107MUM) (**Figure S5)**. Corresponding genes of the oxidoreductase homologs in cluster II are all flanked by a transcriptional regulator (TerR) and a hypothetical protein, but in the case of *A. woodii* and *A. fimetarium*, an additional oleate hydratase is located between TerR and the hypothetical protein (**Figure 4C, Figure S5**).

Moreover, an alignment of all *Acetobacterium* clade 9 homologs (**Figure S6**) revealed that protein VUZ27132.1 contains 26 unique amino acids at position 756–782 which cannot be found in any *Acetobacterium* homolog and only in its two most similar homologs (**Figure 4C, cluster I**). These two homologs (HBC92819.1; WP_190259616.1 **Figure 4C, Figure S5, S6**) belong to *Pelotomaculum* spp. and share ∼77% amino acid sequence identity with protein VUZ27132.1. However, the two *Pelotomaculum* sp. genomes do not encode the complete putative *isr* operon, missing the ferredoxin (ENOG4105DQ9) and the second *hypA*-like homolog (**Figure S5)**. Hence, no corresponding gene of the clade 9 proteins has a ferredoxin from the orthologous group ENOG4105DQ9 located adjacent to it apart from *A. wieringae* ISORED-2 itself and the gene arrangement of the putative *isr* operon is not found in any other organism on NCBI.

### Predicted functional annotation of proteins encoded in the putative *isr* operon

Functional annotation to predict domains and important sites of the proteins encoded by the putative *isr* operon, was performed with InterProScan (48) (**Table S9**). Proteins VUZ27133.1 and VUZ27136.1 were predicted to entirely consist of a HypA-like domain (PF01155, IPR000688), Ni-metallochaperone. Like HypA proteins in other organisms both HypA-like proteins in *A. wieringae* ISORED-2 contain conserved binding properties to Ni^2+^ (backbone amides of residues His2, Glu3) and Zn^2+^ (2 CxxC motifs) (49). Protein VUZ27134.1 belongs to the TIGR00073 family hydrogenase maturation factor HypB and is predicted to also contain a CobW/HypB/UreG, nucleotide-binding domain (PF02492).

The first ∼270 amino acids of protein VUZ27132.1 and protein VUZ27135.1 align to each other (Blastp ID% <26%) but no functional protein domain could be predicted from 1 to 270 amino acids by InterProScan. The remaining part of VUZ27135.1 holds a predicted 4Fe-4S ferredoxin-type, iron-sulphur binding domain (IPR017896) (**Table S10**) containing a 4Fe-4S cluster.

Protein VUZ27132.1 is predicted to have oxidoreductase activity (GO:0016491). According to InterProScan results, protein VUZ27132.1 contains a FAD/NAD(P)-binding domain (IPR023753, PF07992) as well as a dihydropyrimidine dehydrogenase domain II (IPR028261, PF14691) which carries two 4Fe-4S clusters (**Figure 5A**). Moreover, InterProScan results suggest protein VUZ27132.1 contains two additional 4Fe-4S ferredoxin-type iron-sulfur binding domains (PS51379) (**Figure 5A, Table S10**) located at position 289–300 and 848–877 amino acids.

**Figure 5.**
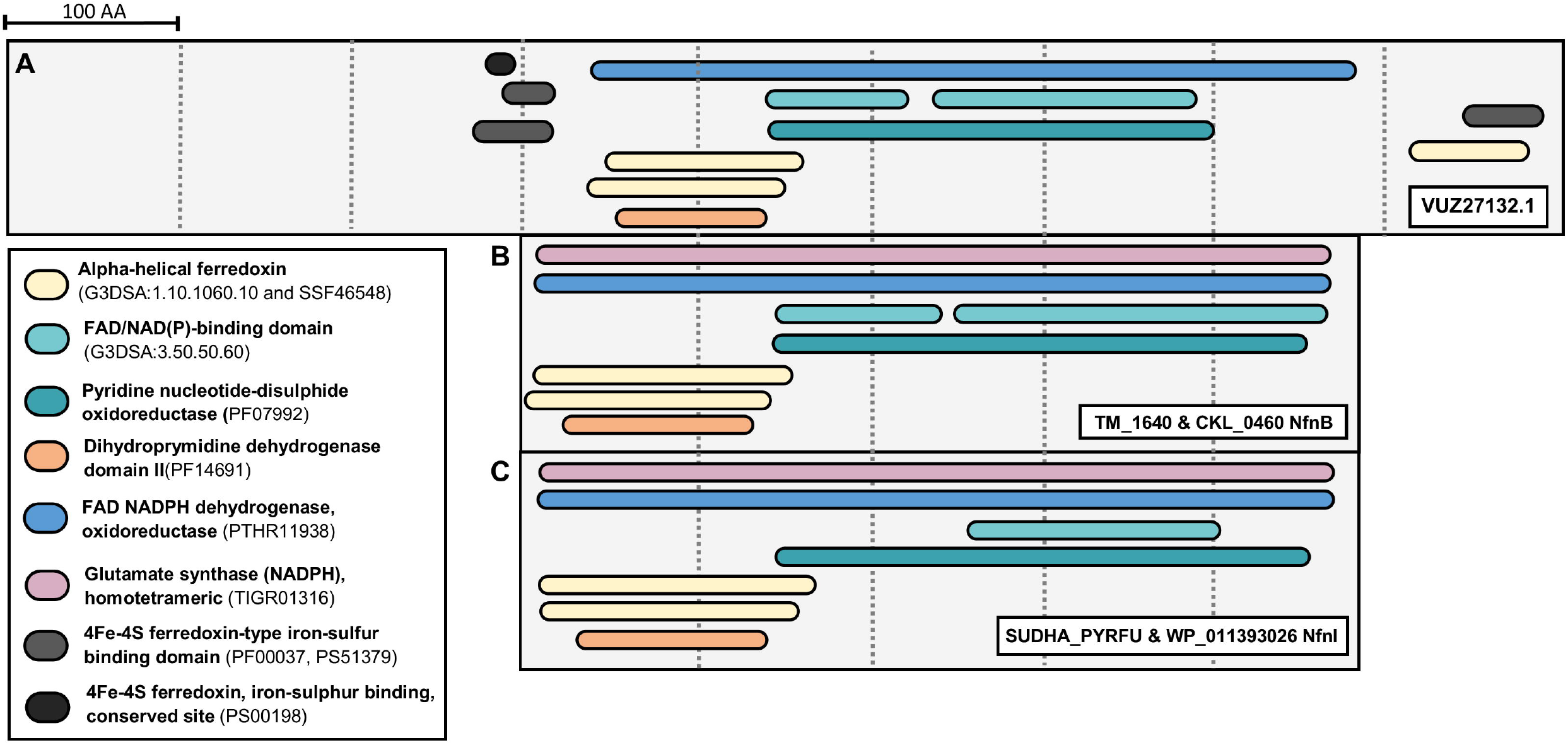
Protein domains of IsrA and the characterized large subunits of Nfn. Results of the InterProScan analysis are shown for each protein (detailed information in **Table S9**).

## Discussion

This study investigated the genetic basis for bacterial isoprene reduction activity previously observed in an isoprene-reducing enrichment (23). The bacteria in this enrichment culture were now identified as a *Comamonas* sp. and a *A. wieringae* sp., named MAG ISORED-1 and ISORED-2 respectively. *A. wieringae* ISORED-2 dominates the isoprene reducing culture (∼89% rel. abundance metagenome sequencing and ∼94% rel. biomass) (**Table 1**) and like other *Acetobacterium* spp. encodes the Wood-Ljungdahl pathway (WLP) for autotrophic growth (50), the Na^+^-translocating ferredoxin: NAD^+^-oxidoreductase (Rnf complex) (51), F_1_F_0_-ATPase (52), the electron transfer flavoproteins (53) and an electron-bifurcating [FeFe]-hydrogenase (54). Even though *A. wieringae* ISORED-2 dominates the isoprene reducing culture a *Comamonas* sp., which shows highest sequence similarity to *Comamonas aquatica* CJG (78.9% ANI, 74.5% AAI) (55), is also present (∼11% rel. abundance metagenome sequencing, ∼6% biomass) (**Table 1**). The involvement of *Comamonas* sp. ISORED-1 in isoprene reduction can therefore not be excluded. However, its relative abundance in H_2_/HCO_3_^-^ /isoprene fed cultures (11%) was lower than in H_2_/HCO_3_^-^ fed cultures (∼23%) based on coverage values from metagenome sequencing (**Table 1**) suggesting that these cells do not benefit from the inclusion of isoprene. Additionally, one of two proteins upregulated upon exposure of *Comamonas* sp. ISORED-1 to isoprene is SpoT (VUZ25726.1), a ppGpp synthetase/hydrolase, indicating that the cells are experiencing nutrient stress. Bacteria respond to nutritional stress by producing (p)ppGpp, which triggers a stringent response resulting in growth arrest and reallocation of cellular resources (33, 56). In *E*.*coli*, fatty acid starvation was found to induce (p)ppGpp accumulation synthesized exclusively by SpoT (57). SpoT interacts with acyl carrier protein (ACP) to likely induce a conformational switch that favours (p)ppGpp synthesis upon fatty acid starvation (58). Interestingly, ACP was one of the proteins downregulated in *Comamonas* sp. ISORED-1 in the presence of isoprene (**Table 1**) and proteins observed in the metaproteome included those for beta-oxidation of fatty acids (**Table S11**). These results are consistent with *Comamonas* sp. ISORED-1 growing on necromass (e.g. fatty acids) and experiencing nutrient stress in the presence of isoprene which slows down cellular growth and metabolism via the (p)ppGpp stringent response.

Isoprene reduction was found to be an induced rather than constitutive trait and comparative proteomics identified 13 proteins upregulated upon isoprene exposure. Apart from *A. wieringae* ISORED-2’s oxidoreductase (VUZ27132.1), no isoprene responsive protein from *A. wieringae* ISORED-2 or *Comamonas* sp. ISORED-1 is predicted by protein function to be involved in redox processes (**Table 2**). This makes the oxidoreductase from the *A. wieringae* lineage, the only likely candidate within the 13 isoprene-responsive proteins that could catalyse the isoprene hydrogenation reaction. The oxidoreductase is encoded in a putative five-gene operon together with the corresponding genes for three nickel-binding chaperones and one 4Fe-4S ferredoxin (**Figure 2A**). Four out of the five proteins encoded in this putative operon were also highly upregulated upon isoprene exposure in *A. wieringae* ISORED-2 (**Figure 1**) and are also found to be unique to the *A. wieringae* ISORED-2 genome by pangenomic comparison of available *Acetobacterium* genomes (**Table S7**). Because closest relative *A. wieringae* DSM 1911 did not exhibit isoprene reducing activity (23), it follows that genes encoding isoprene reduction most likely sit within this unique set. Apart from the corresponding genes for Fe-4S ferredoxin, two HypA-like proteins and the oxidoreductase (VUZ27132.1), no other genes responding to isoprene are unique to the ISORED-2 MAG. Henceforth the operon will be referred to putatively as the isoprene-regulated operon (*isr* operon) and the oxidoreductase (VUZ27132.1) as the putative isoprene reductase or IsrA (gene name *isrA*).

IsrA is predicted to contain a nested FAD and NAD(P)H binding site as well as two pairs of [4Fe-4S]-clusters (**Figure 5A**). The best characterized and only crystallized proteins in the orthologous group of IsrA are the β-subunits of NADH-dependent ferredoxin-NADP^+^-oxidoreductases (Nfn) (**Figure 4AB**, evolutionary group 3). Nfn is an electron-bifurcating enzyme (59), comprising two subunits, NfnA (32.6 kDa) and NfnB (49.8 kDa). Crystal structures from *Thermotoga maritima* (TM_1640) and *Pyrococcus furiosus* (PF1327) revealed that NfnB contains two [4Fe-4S]-clusters as well as binding sites for NADPH and FAD with FAD being the site of electron bifurcation (39, 60, 61) (**Figure 5 BC**). Since IsrA is predicted to contain a FAD/NAD(P)H binding site as well as four 4Fe-4S-clusters and a 4Fe-4S ferredoxin (VUZ27135.1) is encoded in the putative *isr* operon, bifurcation may be a reaction mechanism to contemplate for IsrA. The standard redox potential of the isoprene/methylbutene couple is not known but based on calculation using the estimation of isoprene energy of formation 197 kJ mol^-1^(23, 62, 63) and theoretical stoichiometries with H_2_ as electron donor for the isoprene hydrogenation reaction (62, 64),

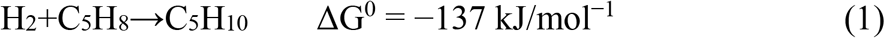

the standard reduction potential (E^0^) of the isoprene/methylbutene couple calculated using Nernst Equation is estimated as follows:

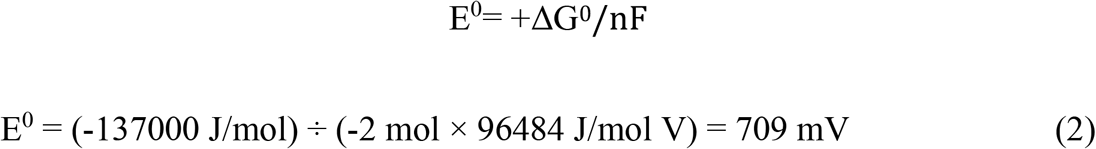

Nernst equation for standard potential of biological systems at pH 7

Eox/red = E^0^ox/red +(2.3 RT/nF) lg [ox]/[red]; 2.3 RT/F= 0.059 at T=289 K, F=96500, R=8.31 Eox/red = E^0^ox/red +(0.059/n) lg [ox]/[red];

Standard electron potential of hydrogen electrode: EH_2_= E^0^H_2_ +(0.059/1) lg(H^+^); E^0^H_2_=0; EH_2_=**-0.059pH**

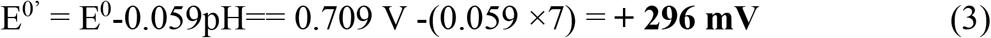

Hypothetically, similar to caffeate reduction, NADH (E^0’^ = -320 mV) derived from the [FeFe]-hydrogenase and Rnf complex could act as the reductant for the exergonic reduction of isoprene to methylbutene (E^0’^ = +296 mV) which is coupled to the endergonic reduction of ferredoxin (E^0’^ = -420 mV) (65). As with caffeate respiration, the reduced ferredoxin could be re-oxidized at the Rnf complex to generate an Na^+^ gradient (32). Out of 12 known flavin-based bifurcating enzymes (59, 65), three are found in *Acetobacterium* spp.: the bifurcating [FeFe]-hydrogenase (54), lactate dehydrogenase/electron transfer flavoprotein (Bf-Ldh) (66) and the caffeyl-CoA reductase (32). However, homology to bifurcating enzymes is not sufficient to guarantee electron bifurcating functionality (59) but as energetics and the binding sites of IsrA support the idea of an electron bifurcating process, it should be considered in future biochemical investigations of IsrA. If isoprene reduction was a linear process, reduction would have to be coupled to ATP synthesis through establishment of an ion gradient, since the reduction of isoprene conserves energy (23).

Beside IsrA, the putative *isr* operon contains three *hyp* genes (**hy**drogenase **p**leiotropic) which encode metallochaperones typically responsible for acquisition and insertion of nickel during maturation of [NiFe]-hydrogenases (67–71). Biosynthesis and maturation of the [NiFe]-hydrogenase active site is a complex multistep process also involving accessory Hyp proteins (HypCDEF) (72). However, the genome of *A. wieringae* ISORED-2 does not harbour *hypCDEF* to produce a functional [NiFe]-hydrogenase. This suggests that HypA and/or HypB might be involved in nickel insertion into an active site of one of the proteins in the *isr* operon. HypA and HypB are known for involvement in the nickel-dependent maturation of other nickel-dependent enzymes besides NiFe hydrogenases e.g. urease in *Helicobacter pylori* (69).

Homologs of IsrA are widely distributed among anaerobic bacteria, but the putative *isr* operon as observed in *A. wieringae* ISORED-2 was not found in any other organism in NCBI. Acquisition of the putative *isr* operon via horizontal gene transfer may be one possible scenario that explains why only *A. wieringae* ISORED-2 harbours the putative *isr* operon. The operon is located in a 44 kbp genomic region containing metabolic genes and is also flanked by mobile genetic elements (**Figure 3**): a *Siphoviridae* provirus and a series of insertion sequences in tandem, which suggests that the putative *isr* operon is placed in a dynamic genomic region of *Acetobacterium wieringae* ISORED-2. Other organisms that also encode the complete putative *isr* operon, from where horizontal gene transfer could have occurred, are yet to be identified.

Homologs of IsrA observed in other *Acetobacterium* spp. share only ∼47–49% amino acid sequence identity and not only are these homologs located in separate subclades (**Figure 4C**) but their corresponding genes are also found in different gene arrangements (**Figure S5**). Together with the inability of other *Acetobacterium* spp. (i.e., *A. woodii* DSM 1030, *A. malicum* DSM 4132 or *A. dehalogenans* DSM 11527) to reduce isoprene as determined experimentally, the phylogenetic analysis provides further evidence that IsrA and its homologs in other *Acetobacterium* spp. have distinct enzymatic functions. Potential enzymatic functions to consider for IsrA homologs are the hydrogenation of unfunctionalized (conjugated) C=C bonds in other unsaturated hydrocarbons found in anoxic environments, such as terpenes, which consist of isoprene building blocks. This might be the case for *Pelotomaculum schinkii* which encodes the most closely related homolog to IsrA (**Figure 4C**). *P. schinkii* is a strictly anaerobic, syntrophic bacterium known to live in electron acceptor depleted environments metabolizing propionate and must resort to using H^+^ and CO_2_ as electron sinks (73). Degradation of propionate is thermodynamically challenging and can only be reached if H_2_ or formate are kept at very low concentration by a syntrophic partner methanogen (73). However, due to its IsrA homolog *P. schinkii* might have the ability to use unfunctionalized (conjugated) C=C bonds in unsaturated hydrocarbons as electron acceptors which could enable them to grow axenically. As a general example, the oxidation of propionate coupled to isoprene reduction would be thermodynamically favourable (23, 62):

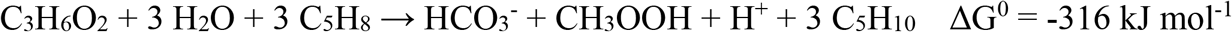

This study provides first evidences for the existence of a putative isoprene reductase. The putative isoprene reductase is of particular interest because of its reduction of a unfunctionalized conjugated C=C bond. IsrA homologs are widespread among various taxonomic groups of strictly and facultatively anaerobic bacteria (*Firmicutes, Spirochaetes, Tenericutes, Actinobacteria, Chloroflexi, Bacteroidetes* and *Proteobacteria*) suggesting that the use of unfunctionalized C=C bonds in unsaturated hydrocarbons as anaerobic electron acceptors is a form of bacterial energy harvesting not previously recognized. While more rigorous physiological/biochemical testing is required to fully understand what the functions of IsrA and its homologs are, the results have both environmental relevance in the context of furthering our understanding of electron sinks in anaerobic environments, and in our understanding of contributing mechanisms to global isoprene turnover.

## Experimental procedures

### Strains and culturing conditions

*Acetobacterium* species *A. woodii* DSM 1030, *A. malicum* DSM 4132, *A. wieringae* DSM 1911 and *A. dehalogenans* DSM 11527 were obtained from Deutsche Sammlung von Mikroorganismen und Zellkulturen (DSMZ, Germany).

Isoprene reducing biomass was grown on H_2_/HCO_3_^-^/(±isoprene) as described previously (23). Isoprene and H_2_ were resupplied every 2 days (RT-PCR; proteomics and cell suspension assays). After 4 (RT-PCR) or 10 (proteomics and cell suspension assays) days of incubation at 30 °C cells were harvested. Isoprene and methylbutene were quantified by gas chromatography using a GasPro PLOT column (60 m x 0.32 mm, Agilent Technologies) as previously described (23).

### Cell suspension assays

Cells from six flasks from H_2_/HCO_3_^-^/isoprene and six flasks from H_2_/HCO_3_^-^ grown cultures were pooled in an anaerobic chamber by pipetting cell aggregates into two separate 6 ml anoxic glass flasks. Cells were washed in minimal media containing 1 mM titanium citrate (23), then OD_600_ (7.5) and volumes (1.57 ml) were adjusted between the two samples. Flasks were crimp sealed and flushed with N_2_ for 30 mins to remove isoprene, methylbutenes and CO_2_. Headspace was measured for isoprene and methylbutene before the experiment was started. H_2_ (0.7 bar), HCO_3_^-^ (60 mM) and isoprene (1 mM) were added, and cells incubated at 30°C (shaking 180 rpm). Headspace (100 μl) was analysed for isoprene depletion and methylbutene production as previously described (23). Liquid samples (0.04 ml) were analysed for acetate. Acetate was analysed as its ethyl ester derivative by GC-FID as previously described (23) but with reduced sample size.

### DNA extraction and Illumina sequencing

DNA was extracted from isoprene reducing cultures anaerobically grown on H_2_/HCO_3_^-^ /(±isoprene) as described previously (23). Libraries were prepared using the Nextera XT DNA Sample Preparation Kit according to manufacturer’s protocol (Illumina). Sequencing reactions were carried out using MiSeq v2 (2×150 bp) chemistry (Illumina) on a MiSeq instrument (Illumina) by the Ramaciotti Centre for Genomics at UNSW (Sydney, Australia).

### RNA extraction and Reverse transcription PCR

Cell aggregates from three flasks were pooled, centrifuged at 10 000 × *g* for 10 min and disrupted in lysis buffer (400 μl) (74) with mechanical agitation (30 Hz for 10 min) in FastPrep Lysis Matrix A tubes (MP Biomedicals). RNA was extracted with sequential phenol-chloroform-isoamyl alcohol (25:24:1) (pH 4.5), 3 M sodium acetate (pH 5.2) and chloroform treatments, precipitated with isopropanol and GlycoBlue™ Coprecipitant (Thermo Fisher Scientific, Australia), resuspended in 35 μl H_2_O and stored at -20 °C. Residual DNA in RNA samples was digested with RNase-free DNase (Qiagen) I and cleaned three times on a Spin Column from PureLink® RNA Mini Kit (Thermo Fisher Scientific, Australia). RNA was quantified with Qubit™ RNA HS Assay Kit (Thermo Fisher Scientific, Australia). RNA samples were stored at -80 °C until use. First strand cDNA was synthesized from 100 ng DNase I-treated total RNA using random hexamer primers from the RevertAid™ First Strand cDNA Synthesis Kit (Thermo Fisher Scientific) following manufacturer’s instruction. In a negative control the M-MuLV reverse transcriptase was replaced with water. Synthesized cDNA was used as template in PCR with the Q5 high-fidelity DNA polymerase (New England BioLabs) using intergenic region primers (**Table S3 & S4**). Chromosomal DNA was used as a template as a positive control (**Figure 2C**).

### Protein extraction and LC-MS/MS analysis

Cells were grown in 8 flasks with H_2_/HCO_3_^-^/isoprene and 8 flasks with H_2_/HCO_3_^-^. To increase cell mass, cells from two flasks where pooled from 8 to 4 samples for each condition, i.e. four replicates for each condition. Cell aggregates were transferred into 2 ml tubes inside the anaerobic chamber, cells were then centrifuged at 10000 × *g* for 10 min and stored at -20°C until use. Harvested cells suspended in 100 μl of lysis buffer (74) were mechanically disrupted in FastPrep Lysis Matrix A tubes (MP Biomedicals) at 30 Hz for 10 min. Crude extracts were passed through a 30 kDa Amicon Ultra-0.5 mL Centrifugal Filters and washed 6 times with 200 μl of 50 mM NH_4_4HCO_3_ buffer (pH 6.9). Protein concentrations were determined with the Quick Start Bradford Protein Assay following manufacturer’s instructions (Bio-Rad Laboratories, Australia), and adjusted to 2 μg μl^-1^ and 10 μl (20 μg) were used for Filter Aided Sample Preparation (FASP) (75–77). Samples were treated with 5 mM dithiotreitol (DTT) at 37 °C for 30 min. Protein lysates were then transferred to 30 kDa Amicon Ultra-0.5 mL Centrifugal Filters and treated following FASP method involving an alkylation step (100 μl of 50 mM iodoacetamide). Trypsin solution (1 μl of a 200 ng μl^-1^) was added for digestions at 37 °C overnight. Peptides were eluted in 2 × 20 μl 50 mM NH_4_HCO_3_ buffer and stored at -20 °C until LC-MS/MS analysis.

Sample analysis was performed at the Bioanalytical Mass Spectroscopy Facility (BMSF) at UNSW. Digested peptides were separated by nanoLC using an Ultimate nanoRSLC UPLC and autosampler system (Dionex, Amsterdam, Netherlands). Samples (2.5 μl) were concentrated and desalted onto a micro C_18_ precolumn (300 μm x 5 mm, Dionex) with H_2_O:acetonitrile (98:2, 0.1% TFA) at 15 μl/min. After a 4 min wash the pre-column was switched (Valco 10 port UPLC valve, Valco, Houston, TX) into line with a fritless nano column (75μ x ∼15 cm) containing C_18_AQ media (1.9μ, 120 Å Dr Maisch, Ammerbuch-Entringen Germany). Peptides were eluted using a linear gradient of H_2_O:CH_3_CN (98:2, 0.1% formic acid) to H_2_O:CH_3_CN (64:36, 0.1% formic acid) at 200 nl/min over 30 min. High voltage (2000 V) was applied to low volume Titanium union (Valco) and the tip positioned ∼ 0.5 cm from the heated capillary (T = 275 °C) of a Orbitrap Fusion Lumos (Thermo Electron, Bremen, Germany) mass spectrometer. Positive ions were generated by electrospray and the Fusion Lumos operated in data dependent acquisition mode (DDA).

A survey scan m/z 350–1750 was acquired in the orbitrap (resolution = 120,000 at m/z 200, with an accumulation target value of 400,000 ions) and lockmass enabled (m/z 445.12003). Data-dependent tandem MS analysis was performed using a top-speed approach (cycle time of 2 s). MS2 spectra were fragmented by HCD (NCE = 30) activation mode, and the ion-trap was selected as the mass analyser. The intensity threshold for fragmentation was set to 25,000. A dynamic exclusion of 20 s was applied with a mass tolerance of 10 ppm.

### Genome assembly and annotation

Quality trimming was performed with BBDuk (http://sourceforge.net/projects/bbmap/). Filtered reads were co-assembled with MegaHIT v1.1.3 (78) and default parameters. Contigs ≥2.5 kbp were manually binned and curated under anvi’o v5.2.0 (79). The contig containing the rRNA operon removed due to its chimeric nature (single chimeric contig detected). MAG completion estimates were obtained with I) anvi’o bacterial SCG profile, II) CheckM v1.1.2 (80) lineage_wf and III) CheckM lineage_wf with domain-specific profiles. Protein coding genes of the metagenome and genomes were predicted with Prodigal v2.6.3 (81).

Metagenome assembled genomes (MAGs) derived from binning were identified and named based on the Genome Taxonomy Database with GTDB-Tk v0.1.3 (82). Predicted proteins were annotated with InterProScan v5.25-64 (83). Results were parsed with a custom script, iprs2anvio.sh (https://github.com/xvazquezc/stuff/blob/master/iprs2anvio.sh) and integrated in the anvi’o workflow. Predicted proteins were assigned to bacterial orthologous groups using the bactNOG database from EggNOG v4.5.1 (84) with EggNOG-mapper v1.0.3-3-g3e22728 (84). Genome annotation was performed with a modified version of Prokka v1.13.3 (85) in which Prodigal generates partial gene calls at the end of contigs in order to minimise differences between Prokka- and anvi’o-based gene predictions.

Prophage/provirus prediction was performed with VirSorter v1.0.6 (86), PHASTER web server (87), Phigaro v2.3.0 (88), and CheckV v0.7.0 with v0.6 database (89).

### Refinement of the operon gene environment

Due to the high coverage of ISORED-2, only reads from the samples containing isoprene were mapped back to the ISORED-2 MAG with Bowtie2 v2.3.4.3 (90), and examined in IGV v2.8.10 (91). The genome assembly graph was visualised with Bandage v0.8.1 (92). ISORED-2 was iteratively reassembled with MIRA v5rc2.

### Pangenome analysis

Pangenomic analysis of the genus *Acetobacterium* with eight reference *Acetobacterium* genomes (**Table S5**) was performed with anvi’o v5.2.0 (79) following the standard pangenomics workflow (http://merenlab.org/2016/11/08/pangenomics-v2). Genes were clustered with MCL inflation values of 2, 4 and 6 (93). Gene clusters were grouped based on their presence in all 9 genomes (core), at least 4 out of 9 genomes (soft core) or unique to the organism (singleton). Average Nucleotide Identity (ANI) between *Acetobacterium* genomes was calculated with pyani v0.2.7 (94). Average Amino Acid Identity was calculated with CompareM v0.0.23 (https://github.com/dparks1134/CompareM).

### MS data analysis

The raw MS data was processed using MaxQuant software (version 1.6.2.1) (95) and searched against a custom database of all predicted proteins in the metagenome of the isoprene reducing culture (6517 sequences). Enzyme specificity was set to trypsin/P, cleaving C terminal to lysine and arginine and a maximum number of two missed cleavages allowed. Carbamidomethylation of cysteine was set as a fixed modification and oxidation of methionines, acetylation of protein N termini were set as variable modifications. The minimum peptide length was set to 7 amino acids and a maximum peptide mass was 4600 Da. The minimal score for modified peptides was 40 and the minimal delta score for modified peptides was 6. Peptide intensities were normalised using MaxLFQ (96). Downstream analysis was performed in R v3.5.1 with the package DEP v1.4.0 (97). First, MaxQuant output data was filtered, retaining only proteins detected by at least two unique peptides, and detected in all replicates. In order to reduce the influence of the changing community composition and their relative contributions towards the total metaproteomic data, the metaproteomic data was partitioned based on the source MAG and analysed separately in DEP as follows. LFQ intensities were normalised with vsn (98) and missing values imputed by left-censored imputation (MinProb function). Differential expression analysis was conducted with limma (99). Proteins were considered differentially expressed if they had an adjusted (FDR) *p-*value ≤ 0.05, and a log2 fold change (LFC) ≥ 2 or ≤ -2.

The number of peptide spectrum matches per protein were used to quantify the biomass contribution of each organism to the community (100).

### Phylogenetic analyses

#### Molybdopterin oxidoreductase VUZ27132.1 (putative isoprene reductase)

Phylogenetic trees were constructed based on all protein sequences from the EggNOG database v4.5.1 (84) matching the orthologous group of VUZ27132.1 (ENOG4107QZ5), and its archaeal homolog (arCOG01292). Additional ENOG4107QZ5 sequences from other *Acetobacterium* genomes and characterized enzymes from the literature were included. An additional 1000 top hit sequences to VUZ27132.1 retrieved from NCBI (blastp search on 10 Dec 2018) were clustered with CD-HIT v4.6 (-s 0.8 -c 0.8) and added to the dataset (101). All sequences were aligned using MAFFT v7.313 (mafft-linsi) (102). Alignment was manually trimmed to restrict the phylogeny to the core/conserved region of the proteins, equivalent to positions 158–779 of 901 residues in VUZ27132.1. In addition, gap-rich columns were removed from the manually-trimmed alignment with BMGE v1.12 (-m BLOSUM30 -g 0.9 -h 1) (103). The phylogenetic protein tree was constructed with IQ-TREE v1.6.7 (104) using a LG+I+G4 model (105) and 10000 ultrafast bootstrap replicates (106). Trees were visualized using iTol interactive tree of life https://itol.embl.de/tree/4918010318257311557452600 (https://itol.embl.de/).

#### Clade 9 phylogeny

Additional top 1000 records from NCBI were retrieved on 16 April 2021 to recover recently deposited IsrA (VUZ27132.1) orthologs. Only sequences with ≥50% identity and ≥75% coverage were added to the original dataset and identical sequences removed. Orthologs from recently sequenced *Acetobacterium* genomes were also included (107). A total of 1063 sequences were aligned with MAFFT-L-INS-i v7.407 (108). Resulting alignment was trimmed in BMGE v1.2 with permissive options (-m BLOSUM30 -g 0.9 -h 0.9) (103). The tree was inferred with IQ-TREE v2.1.2 (109) under EX_EHO+R10 substitution model (110) and 1000 ultrafast bootstrap replicates with nearest neighbour interchange optimisation (--bnni) (111).

Gene neighbourhoods of clade 9 proteins were examined with EFI-GNT (112) (search performed on 18 June 2021).

## Data availability

Raw sequencing data and annotated MAGs have been deposited in ENA under project PRJEB30289 (ERP112722). Metaproteomic data is available at the PRIDE database (PXD023683).

## Acknowledgements

We thank Ling Zhong for her assistance with mass spectrometry in BMSF-UNSW and for her guidance and help with proteomic analysis. We also thank Gene Hart-Smith for his help with the metaproteomic data analysis. Furthermore, we thank Daniel E. Ross for kindly providing the amino acid sequences of *A. malicum* DSM 4132 prior to publication. MK and ML were supported by an Australia-India Strategic Research Fund grant (AISRF48508). MRW and XVC acknowledge support from the New South Wales State Government RAAP scheme and the UNSW RIS scheme.

