## Supplementary Figure for "Evidence for a putative isoprene reductase in *Acetobacterium wieringae*"

### Isoprene reduction is induced in the presence of isoprene, $H_2$ and $HCO_3^-$

To test whether isoprene reduction in an *Acetobacterium*-dominated (rel. abundance 16S rRNA gene amplicon sequencing 92–100%) homoacetogenic enrichment culture (1) is constitutive or inducible, cell suspensions of cells pre-grown with $H_2/HCO_3^-$ or $H_2/HCO_3^-$/isoprene were prepared in phosphate buffered minimal media. Cell suspensions ($OD_{600}$ 7.5 in each flask) were incubated with $H_2$, $HCO_3^-$ and isoprene and monitored for isoprene consumption and production of methylbutene, acetate and formate.

In cell suspensions containing cells pre-grown with $H_2/HCO_3^-$ (**Figure S1A**), acetogenesis commenced immediately at 70 nmol $min^{-1}$ for 135 min and then 24 nmol $min^{-1}$ thereafter. A 100 min lag in isoprene reduction was observed, after which methylbutene formation commenced at 1.25 nmol $min^{-1}$. The lag phase in isoprene reduction indicates that there is an induction process and may reflect the time required for induction of the enzymes catalyzing isoprene hydrogenation. In contrast, in cell suspensions containing cells pre-grown with $H_2/HCO_3^-$/isoprene (**Figure S1B**), both isoprene reduction and acetogenesis commenced immediately. Initial methylbutene formation and acetogenic rates were 30 nmols $min^{-1}$ and 37 nmols $min^{-1}$, respectively. Isoprene reduction stopped after 135 mins, at which time the acetogenic rate increased to 77 nmol $min^{-1}$.

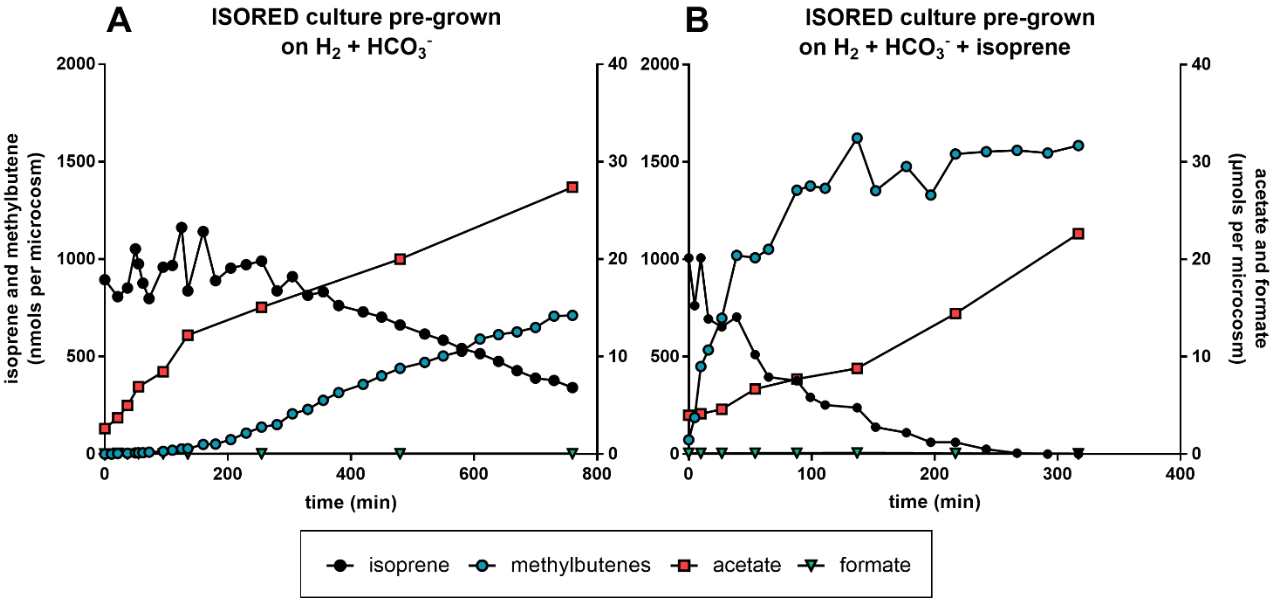


**Figure S1 Induction of isoprene reduction during acetogenesis from $H_2$ plus $CO_2$ by isoprene reducing culture dominated by *A. wieringae*.** Cell suspensions of enrichment culture pre-grown on $H_2/HCO_3^-$ without (**A**) and with isoprene (**B**) were incubated shaking under $N_2$ atmosphere at 30 °C in the presence of 0.5 bar $H_2$, 40 mM $HCO_3^-$ and 1 mM isoprene. Note that the time scales between A and B are different and that on the left y-axis the unit is nmols per microcosm and on the right y-axis it is µmol per microcosm. Note this data results from a single, representative experiment and has no replicates.

### Promotor prediction isoprene-regulated operon

**DNA sequence analysis.** Promoter prediction was performed using the BPROM program and operon prediction was performed with FGENESB which are available through the Softberry website (www.softberry.com).

For promoter-prediction analysis (BPROM) revealed a potential transcription start site around 44 bp upstream of the open reading frame (ORF) 1 (ISORED2_03545; putative isoprene reductase) start codon (sequence of -10 box 'TGTTATAAT' and sequence of -35 box 'ATGTCA') (**Figure S2**). Two potential transcription-factor (RNA polymerase sigma factor rpoD17 and ihf) binding sites were predicted at 58 bp and 38 bp upstream of the ORF 1 start codon. Although a strong candidate, this transcription start site has not been verified by 5'RACE or equivalent. Additionally, 4 transcription factor (TF) binding sites could be identified 13 bp upstream of the ISORED2_03545 start codon (using FIMO with the CollectTF database, output filtered by p-value (<=0.0001), q-values (<=0.05), keeping only matches in the forward sequence). They are highly suggestive binding sites for a Fur (ferric uptake regulator) or NikR (nickel uptake regulator) type of TF, belonging to COG0735 and COG0864 respectively.

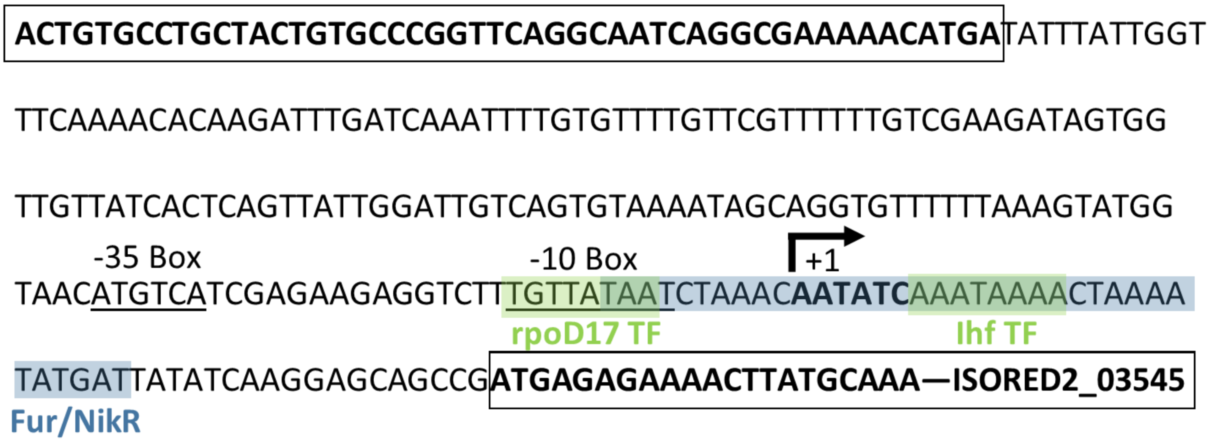


**Figure S2 Promoter region of isoprene operon.** Prediction analysis (BPROM) revealed a potential transcription start site around 44 bp upstream of the open reading frame (ORF) (ISORED2_03545) start codon. Two potential transcription-factor (RNA polymerase sigma factor rpoD17 and ihf) binding sites were predicted at 58 bp and 38 bp upstream of the ORF 1 start codon. Additionally, 4 transcription factor (TF) binding sites could be identified 13 bp upstream of the ISORED2_03545 start codon. They are highly suggestive binding sites for a Fur (ferric uptake regulator) or NikR (nickel uptake regulator) type of TF.

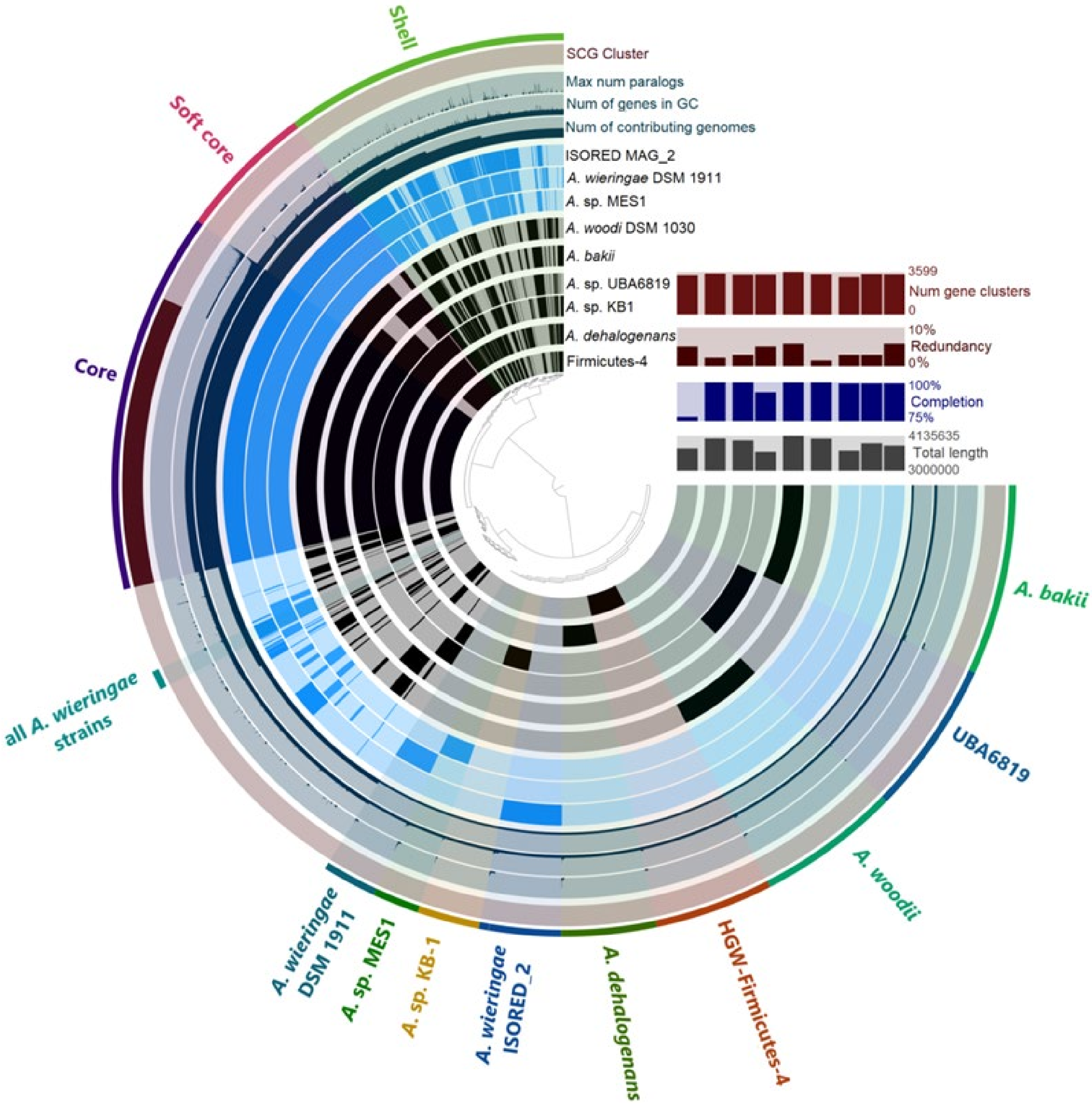


**Figure S3 Pangenome analysis of nine *Acetobacterium* genomes.** List of complete *Acetobacterium* genomes used for pangenome analysis with anvi'o (**Table S5**). Each of the 8,190 gene clusters contains one or more genes contributed by one or more isolate genomes. The "core" selection corresponds to the gene clusters that contain genes from all the genomes. The "soft core" selection corresponds to gene clusters that contain genes from at least 7 genomes and the shell from at least 4 genomes. "Singletons" selection corresponds to clusters that contain one or multiple genes from a single genome. Genes unique to the *A. wieringae* ISORED-2 MAG and other *Acetobacterium* genomes or MAGs are shown (Supplement **Table S6**). In-set bar graphs show the number of gene clusters, percentage genome completion, percentage redundancy and total length for each lineage.

### Genome environment of the putative isoprene-regulated operon

The putative isoprene-regulated operon is located between 69,745-75,048 bp in a 90,374 bp contig (**Figure 3B**). The first half of this contig contains mainly protein-coding genes of viral origin (exact coordinates depend on the prediction tool). The mean contig coverage and the coverage of the proviral portion are close to the values for the MAG indicating no active viral replication. The provirus (*Siphoviridae*) shows an average amino acid identity of 59.11% with the *Erysipelothrix* phage Φ1605 (2) based on CheckV, and tBLASTx of many of the viral proteins also show similar identities with several *Streptococcus* phages recently sequenced (3). This proviral region also appears in other *Acetobacterium* spp. genomes including *Acetobacterium wieringae* DSM 1911 and *Acetobacterium* sp. KB-1 (**Figure 3AC**).

The contig contains three different Ser-recombinases (integrases). Two of them, the ones adjacent to the provirus (**Figure 3**), show very high identity values with recombinases found in *A. wieringae* DSM 1911, *A.* sp. MES1, and *A.* sp. KB-1 and might be part of the provirus itself.

As the provirus and recombinases were nearly the only sequences sharing some resemblance to extant assembled *Acetobacterium* spp. genomes, the contig encoding the putative isoprene-regulated operon was carefully examined: I) by evaluating the placement of the contig in the assembly graph (possible mis-binning); and II) by mapping the reads back to the MAG to assess the possibility of misassemblies. Examination of the metagenome assembly graph in bandage clearly shows two main subassemblies in the MegaHIT assembly, corresponding to the genomes of the two MAGs recovered through binning. Visualisation in IGV provided evidence of a potential misassembly around 16,600-16,750 bp based on a severe drop in the coverage (~1% of mean coverage). Therefore, ISORED-2 was reassembled iteratively using MIRA v5rc2 in a similar way as described by (4).

After 6 iterations, the reassembled ISORED-2 showed a size increase of ~40 kbp (only contigs >2.5 kbp), but more importantly, a more polished isoprene-regulated operon contig. The polished contig dropped the first 16,660 bp (at the point of the coverage drop) but increased its total length to 178,423 bp. This contig showed a higher degree of synteny with the genome of *Acetobacterium* sp. KB-1 (**Figure 3C**), the most complete *Acetobacterium* spp. genome to date and the only one assembled in a single contig. While MIRA was able to extend the putative isoprene-regulated operon contig and provide a better overlook of the gene neighbourhood of the operon, it did it at the expense of collapsing a number of insertion sequences (IS) present around what would be the end of the original contig after the last Ser-recombinase gene, around ~73,550 and 80,500 bp. Most of this IS-rich region have coverages between 200-300x, with exception of the IS21 between 76,632-79,180 bp that has a coverage 500-750x. With most of the contig with a maximum coverage of 100-150x (only isoprene sample mapped), it is fair to assume that all, if not most, IS present here appear in tandem repeats.

**Figure S4 Full uncollapsed clade 9 phylogenetic tree (next page).** Detailed version of Figure 4C showing all sequences included in the analysis. Protein sequences from *Acetobacterium* spp. are highlighted in magenta. Circles at internal nodes indicate ultrafast bootstrap support values. Coloured circles at tip nodes indicate phylum.

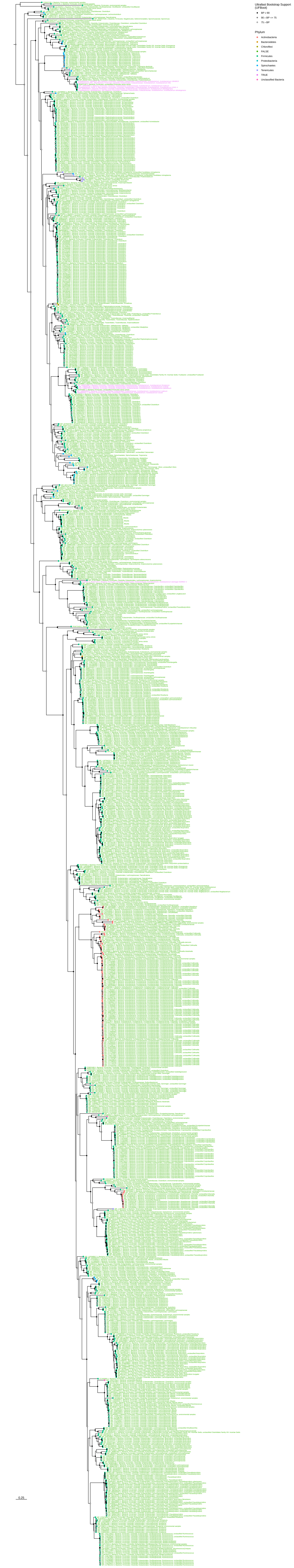
Ultrafast Bootstrap Support (UFBoot)
BP ≥ 90
90 > BP => 75
75 > BP
Phylum
Actinobacteria
Bacteroidetes
Chloroflexi
FALSE
Firmicutes
Proteobacteria
Spirochaetes
Tenericutes
TRUE
Unclassified Bacteria
0.25

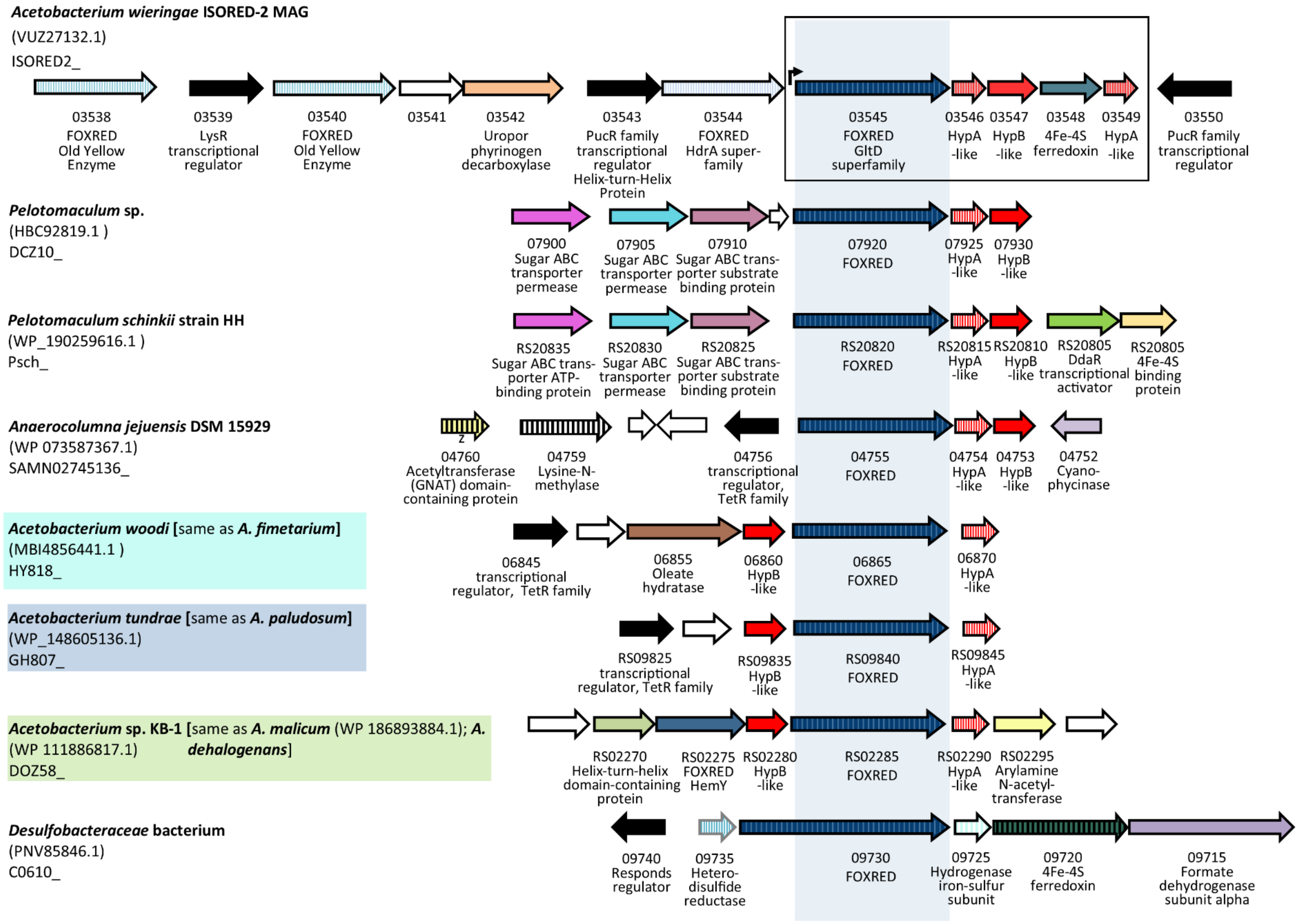


**Figure S5 Gene arrangements of isoprene upregulated putative oxidoreductase IsrA from the *A.wieringae* ISORED-2 MAG and selected clade 9 homologs.** Clade 9 proteins from two *Pelotomaculum* sp. are most closely related to IsrA. The gene environment of clade 9 homologs in other *Acetobacterium* spp. is different to *A. wieringae* ISORED-2. *Acetobacterium* spp. gene arrangements from cluster III (green underline; *A.* sp KB-1; *A. malicum* (WP 186893884.1); *A. dehalogenans* **Figure 4C**) are identical. Gene arrangements in *Acetobacterium* spp. from cluster II (blue and turquoise underline **Figure 4C**) are very similar except that *A.woodi* and *A.fimetarium* also encode an Oleate hydratase and *A. tundrae* and *A. paludosum* do not. *Desulfobacteraceae* and *Syntrophobacteraceae* contain a longer homolog (~1,300-1,400 AA) which is not located next to *hypA* or *hypB*. The *Desulfobacteraceae* bacterium is shown as an example how these gene arrangments are organized. Blank arrows indicated proteins of unknown function.

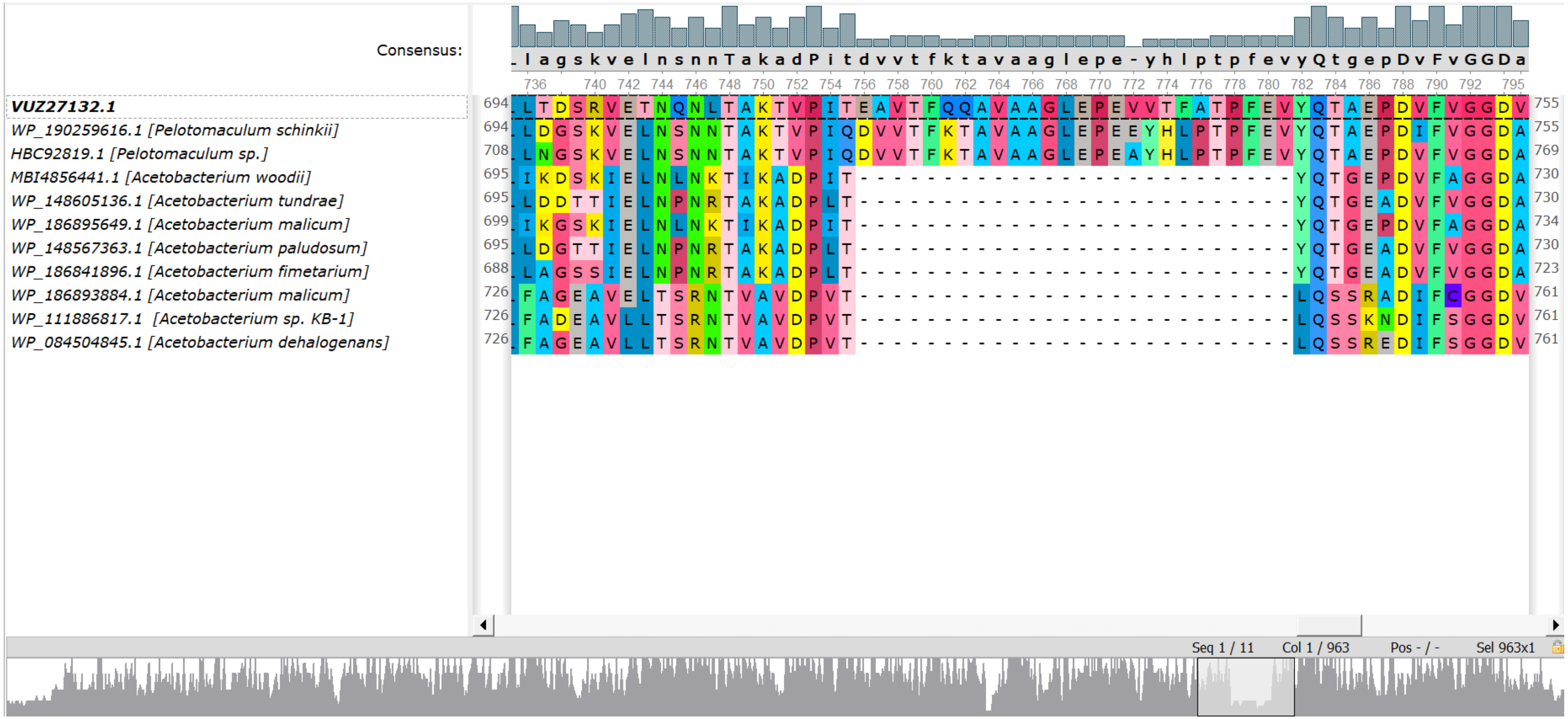


**Figure S6 Amino acids alignment of IsrA (VUZ27132.1), its closest homologs in *Pelotomaculum* sp. and clade 9 homologs in other *Acetobacterium* spp.** Only amino acid positions 735-796 are shown. IsrA contains 26 unique amino acids that cannot be found in other *Acetobacterium* spp. and only in two clade 9 homologs most closely related to protein VUZ27132.1 which belong to *Pelotomaculum* sp. (**Figure 4C**). The sequences were aligned with MAFFT-L-INS-i v7.407. The colors indicate amino acids of different biochemical properties as displayed in UGENE.
